# Variation of the meiotic recombination landscape and properties over a broad evolutionary distance in yeasts

**DOI:** 10.1101/117895

**Authors:** Christian Brion, Sylvain Legrand, Jackson Peter, Claudia Caradec, David Pflieger, Jing Hou, Anne Friedrich, Bertrand Llorente, Joseph Schacherer

**Affiliations:** Université de Strasbourg, CNRS, GMGM UMR 7156, F-67000, Strasbourg, France; CNRS UMR7258, INSERM U1068, Aix Marseille Université UM105, Institut Paoli-Calmettes, CRCM, Marseille, France

**Keywords:** recombination, meiosis, loss of heterozygosity, return to growth, protoploid yeast

## Abstract

Meiotic recombination is a major factor of genome evolution, deeply characterized in only a few model species, notably the yeast *Saccharomyces cerevisiae.* Consequently, little is known about variations of its properties across species. In this respect, we explored the recombination landscape of *Lachancea kluyveri,* a protoploid yeast species that diverged from the *Saccharomyces* genus more than 100 million years ago and we found striking differences with *S. cerevisiae.* These variations include a lower recombination rate, a higher frequency of chromosomes segregating without any crossover and the absence of recombination on the chromosome arm containing the sex locus. In addition, although well conserved within the *Saccharomyces* clade, the *S. cerevisiae* recombination hotspots are not conserved over a broader evolutionary distance. Finally and strikingly, we found evidence of frequent reversion of meiotic commitment to mitotic growth allowing allele shuffling without meiosis completion. Identification of this major but underestimated evolutionary phenomenon illustrates the relevance of exploring non-model species.

**Author summary:** Meiotic recombination promotes accurate chromosome segregation and genetic diversity. To date, the mechanisms and rules lying behind recombination were dissected using model organisms such as the budding yeast *Saccharomyces cerevisiae*. To assess the conservation and variation of this process over a broad evolutionary distance, we explored the meiotic recombination landscape in *Lachancea kluyveri,* a budding yeast species that diverged from *S. cerevisiae* more than 100 million years ago. The meiotic recombination map we generated revealed that the meiotic recombination landscape and properties significantly vary across distantly related yeast species, supporting that recombination hotspots conservation across yeast species is likely associated to the conservation of synteny. Finally, the frequent meiotic reversions we observed led us to re-evaluate their evolutionary importance.

## Introduction

Accurate chromosome segregation at the first meiotic division often requires crossovers (COs) between homologous chromosomes [1]. Meiotic COs result from the repair of programmed DNA double strand breaks (DSBs) by homologous recombination, that inherently yields gene conversions (GCs) [2]. Only a fraction of DSBs yields CO-associated GCs, the remaining yielding GCs without associated COs, called non-crossovers (NCOs). Overall, meiotic recombination provides a nearly limitless source of genetic diversity by rearranging allelic combinations and therefore plays a key role in evolution.

The budding yeast *S. cerevisiae* has long been a model to dissect recombination mechanisms. Meiotic recombination is initiated by Spo11-catalyzed DSBs [3] within nucleosome depleted regions enriched in gene promoters [4] in the context of a highly ordered chromatin structure [5]. Meiotic DSBs and COs occur within well-characterized hotspots across the genome. The mapping of 4,163 COs and 2,126 NCOs from 46 *S. cerevisiae* tetrads [6] showed that the frequency of meiotic recombination events correlates with previously localized DSBs [7,8]. It was also highlighted that the frequency of segregating chromosome without a CO is low in *S. cerevisiae* (1 out of 46 tetrads), supporting the importance of at least one CO per chromosome for accurate segregation. This obligatory CO [9,10] might be a consequence of CO interference [11].

Meiotic DSBs and associated recombination hotspots have long been expected to be extremely unstable across evolution because of their supposed inherent self-destructive nature [12]. This is the case in some mammalian species where DSB hotspots are determined by the sequence specific DNA binding of the fast evolving PRDM9 methyl transferase protein [13]. Remarkably, meiotic recombination initiation hotspots show a good conservation within *Saccharomyces* species despite a sequence divergence almost 10-fold higher than between human and chimpanzees that completely lack hotspot conservation [14–17]. The same observation was made for two *Schizosaccharomyces* species [18]. This likely results from the fact that DSB hotspots in yeasts lie in gene promoters, which are conserved functional elements and do not depend on sequence specific elements such as PRDM9 [15,19]. However, the *Saccharomyces* species have almost completely collinear genomes, which could favor hotspot conservation.

Here, we sought to assess the variation of the meiotic recombination landscape as well as properties by taking advantage of the broad evolutionary range of the Saccharomycotina species [20]. We explored the recombination landscape of the protoploid *Lachancea kluyveri* yeast species (formerly *Saccharomyces kluyveri)* that shares the same life cycle as *S. cerevisiae* but is a distantly related species, which diverged from it prior to a whole-genome duplication event, *i.e.* more than ~100 million years [21,22].

## Results and discussion

### *L. kuyveri* meiotic recombination map reveals a high frequency of 4:0 segregating regions

To explore the *L. kluyveri* meiotic recombination landscape genome-wide we generated a diploid hybrid by mating two polymorphic haploid isolates, NBRC10955 (*MATa*) and 67-588 (*MATα*) [23]. The NBRC10955/67-588 hybrid contains approximately 85,705 polymorphic sites distributed across the eight homologous chromosomes, resulting in a genetic divergence of ~0.7%. It sporulates efficiently, with a spore viability of 73% (Figure S1). Overall, the divergence is in the same order of magnitude to those of the S288c/SK1 and S288c/YJM789 hybrids used for meiotic tetrad analysis in *S. cerevisiae* [6,24].

By whole genome sequencing, we genotyped all four viable spores from 49 meioses of the NBRC10955/67-588 hybrid. We selected a set of 56,612 reliable SNPs as genetic markers to define the allelic origin along the chromosomes (see material and methods and Figure S2). This resulted in a median distance of 196 bp between consecutive markers, with only 27 inter-marker distances higher than 5 kb. These markers are well distributed across the genome, with the exception of the rDNA array on chromosome H and the subtelomeric regions. Subtelomeric regions carried low quality sequences and translocation events, which were manually determined. To avoid spurious identification of recombination events at the ends of the chromosomes, we disregarded some subtelomeric regions, adapting case by case their size according to the presence of translocations and/or to the poor sequence quality (Figure S3). Altogether these regions range from 2 to 32 kb in size and represent 2% of the entire genome (251 kb).

From these data, we performed a detailed analysis of individual recombination events using the CrossOver program [25]. Across the 49 meioses, we identified 274 NCOs and 875 single COs (Figure 1A), 541 of which were associated with a detectable GC tract. Small GC tracks associated or not to COs might not overlap with enough genetic markers and the corresponding number could be underestimated. Surprisingly, we observed a high frequency of double COs, which correspond to two COs at the same location involving the four chromatins (138 events, 16% of the total COs) (Figure 1A) as well as a high frequency of regions with a 4:0 allelic segregation (505 regions) (Figure 1A). Previous *S. cerevisiae* studies reported mostly single COs and GC tracts with 3:1 allelic segregation [6,24,25]. These types of events involving the four chromatids simultaneously are usually expected to reflect mitotic recombination events preceding meiotic entry and are generally disregarded [6,24]. In our case, their quantitative importance prompted us to examine them more carefully.

**Figure 1.**
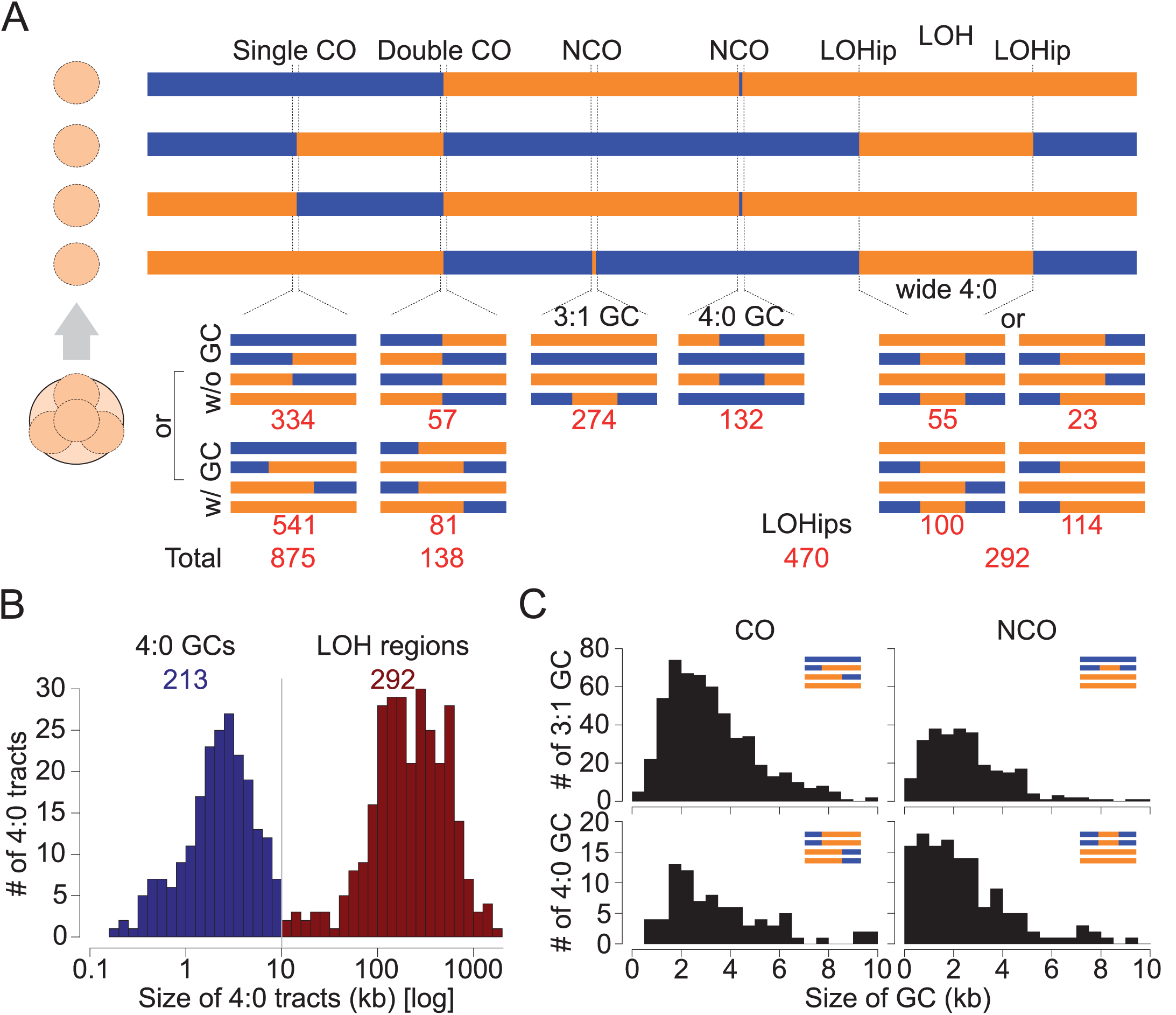
*L. kluyveri* recombination events classes. (A) Allelic segregation patterns observed within a four-spore viable tetrad. CO: crossover, NCO: non-crossover, GC: gene conversion, LOH: loss of heterozygosity, LOHip: LOH initiation point. Red values correspond to the number of instances detected across the 49 tetrads studied. (B) The 4:0 tracts were distinguished according to their sizes. The threshold of 10 kb was defined using the end of the peak corresponding to the short 4:0 tracts that are thought to result from meiotic 3:1 GCs after RTG. (C) Comparison of the distribution of GCs sizes. Top left: 3:1 GCs associated to COs. The median size is 2.8 kb. Bottom left: 4:0 GCs associated to COs. The median size is 3.0 kb. Top right 3:1 GCs associated to NCOs. The median size is 2.3 kb. Bottom right: 4:0 GCs associated to NCOs. The median size is 1.9 kb.

### High meiotic reversion frequency leads to extensive LOH events in meiotic products

We identified a total of 505 regions with a 4:0 segregation. The size distribution of these tracts follows a bimodal pattern with an almost equal partitioning of events in both clusters (Figure 1B). Based on this distribution, we defined two categories: short 4:0 regions (median = 2.3 kb, N = 213) and large 4:0 regions (median = 224 kb, N = 292) with a size cut-off of 10 kb (Figure 1B). The size distributions of the 213 short tracts smaller than 10 kb associated with COs and NCOs mimic the size distributions of the 3:1 conversion tracts associated with COs and NCOs, respectively (Figure 1C). The 292 large regions with a 4:0 segregation bear all the characteristics for being the results of mitotic G2 COs prior entry into meiosis that lead to large regions of loss of heterozygosity (LOH), ranging from 11 to 1,882 kb.

These 4:0 segregating regions and the double COs were found within 38 tetrads. In these tetrads, the number of large 4:0 regions varies between 2 to 18 and their cumulative size ranges from 5% to 63% of the genome (Figure 2A and B). All these large LOH regions are unique to each tetrad and the conserved allelic version seems random leading to an equal frequency of each parental allele (Figure 2B). However, these large LOH events are not equally distributed along the genome. Centromere-proximal regions show a very low rate of LOH events. Every chromosome carries LOH events with the exception of the 1-Mb introgressed region (Sakl0C-left) carrying the *MAT* locus that displays a systematic 2:2 segregation (Figure 2).

**Figure 2.**
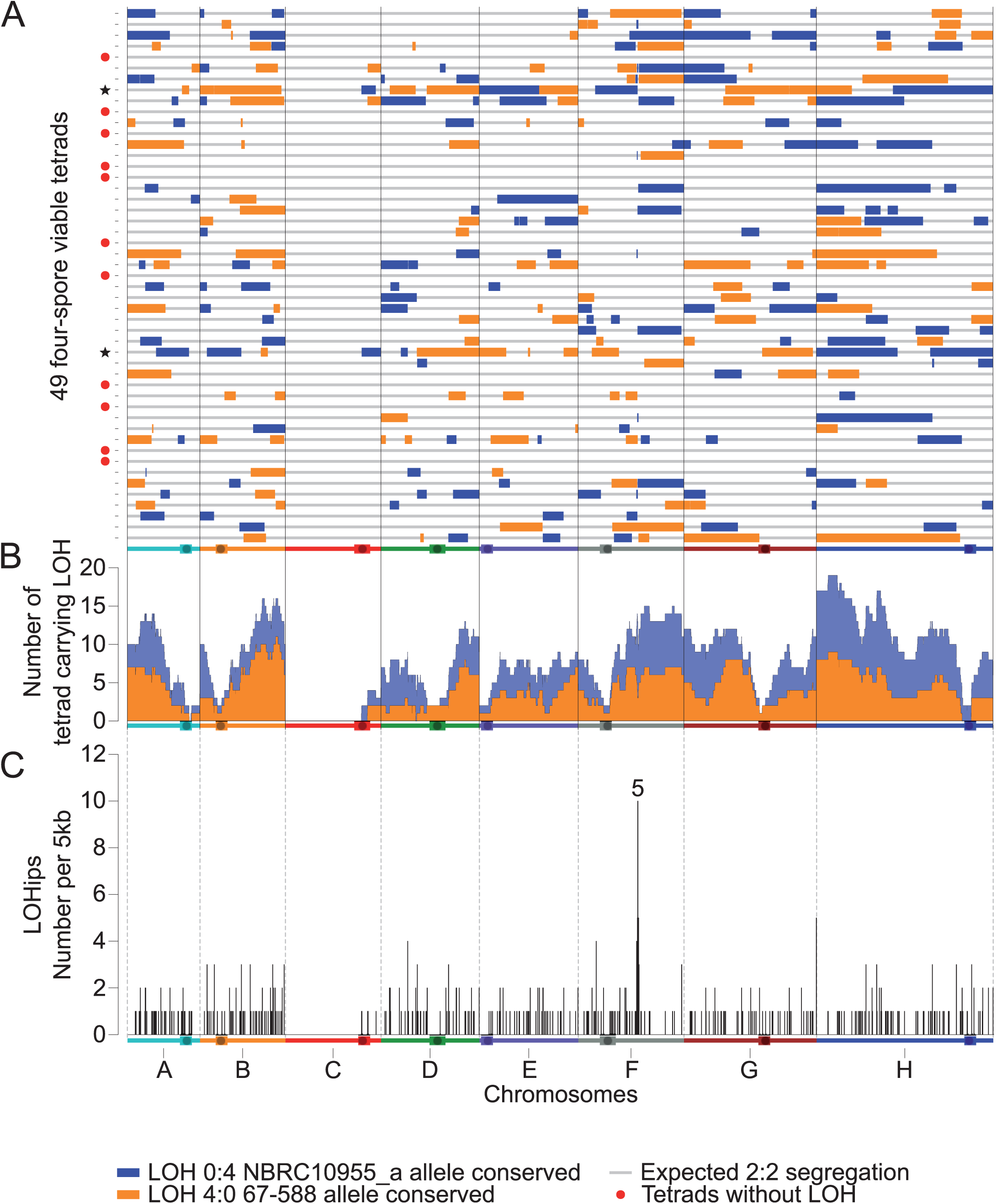
Distribution of LOH events across the 49 sequenced tetrads. (A) Location of the LOH regions along the genome for each tetrad. Colors correspond to the allelic version conserved. Tetrads flagged by a red dot do not carry any LOH. The two tetrads flagged by a black star have all their centromeres included in LOH regions. (B) Frequency of LOH events along the genome across the 49 tetrads using a 5kb sliding window. The frequency of LOH conserving NBRC10955 alleles (blue area) is added to the frequency of LOH conserving 67-588 alleles (orange area) to provide the total frequency of LOH events along the genome. (C) Frequency of LOHips along the genome using a 5 kb window. Region 5 is a hotspot of LOHips and overlaps with the *VMA1* gene. A total of 470 LOHips were detected across the 38 tetrads with LOH.

Overall, the frequency and repartition of 4:0 segregating regions and double COs support a high proportion of mitotic events prior meiotic entry. Such a mitotic genomic instability is unexpected, except from some mutants defective in cell-cycle checkpoints for instance. To test the mitotic stability of the NBRC10955/67-588 hybrid, we induced an accumulation of mitoses by generating two independent mutation accumulation lines. They went through 12 single cell bottlenecks on rich medium representing ~ 240 generations. The sequence analysis of the two evolved diploids as well as the initial parental diploid (only ~ 40 generations) revealed no LOH event (Figure S4). In conclusion, the frequency of mitotic recombination on rich medium of the NBRC10955/67-588 hybrid is too low to explain the high amount of LOH events in the dissected tetrads, and therefore that these events most likely occurred during meiosis.

The only alternative explanation left is that the high frequencies of observed LOH, double CO and 4:0 GC events are due to the so-called “return to mitotic growth” (RTG). RTG is a meiotic reversion where cells enter into meiotic prophase, initiate inter-homolog meiotic recombination, but return to vegetative growth before meiosis I [26,27]. The resulting recombination events are indistinguishable from inter-homolog mitotic G2 events but come from the abortive induction of the meiotic program (Figure S5). In support of a meiotic origin, we found a lack of LOH events near the centromeres, similar to canonical meiotic recombination events. However, two out of the 38 tetrads display LOH regions including all of the eight centromeres. A possible explanation for these two cases would be that RTG occurred after meiotic anaphase I. Such events would have escaped the meiotic commitment thought to be irreversible after meiotic prophase in *S. cerevisiae* through a positive feedback expression of the Ndt80 meiotic transcription factor [28].

Finally, RTG implies at least some residual growth on sporulation medium. We therefore isolated individual NBRC10955/67-588 hybrid cells on sporulation medium, and after two days, we did observe small colonies of about 0.5 mm of diameter, corresponding approximately to ten generations. This indicates that potassium acetate used as sporulation medium and cellular nutrients storage are enough to ensure residual growth, before starting sporulation. A possible explanation is that potassium acetate does not induce a strong enough starvation stress, due to the more respiratory lifestyle of this species [29,30].

Since RTG was highlighted, it has been suspected to happen in natural conditions and induce phenotypic diversity by generating new allelic combinations [31]. The ease with which *L. kluyveri* undergoes RTG leads us to consider that its frequency in natural conditions is high. The resulting LOH regions allow the expression of recessive alleles and new combination of interactive alleles. A recent study also demonstrated that COs induced during RTG can be used to generate an allelic reshuffled population allowing genetic linkage analysis [31]. Interestingly, RTG is initiated in stress conditions and the resulting allelic shuffling might increase the fitness to environment, immediately assuring cell survival. Therefore, the fact that RTG is easily induced in some yeast species can reveal a new mechanism of adaptation to stress, the generality of which remaining to be determined.

### *VMA1* corresponds to a meiotic hotspot for double COs and LOHs

Our recombination map revealed that the main hotspot associated with RTG events, *i.e.* double COs, 4:0 NCOs and LOH regions corresponds to the *VMA1* gene located on chromosome F (hotspot 5, Figure 2C and Figure S6). This gene encodes a subunit of a vacuolar ATPase and the PI-Sce1 endonuclease (also called Vde1) in *S. cerevisiae* as well as in *L. kluyveri* [32,33]. This is a site-specific endonuclease active during meiosis that promotes a homing reaction (Figure S7). In a heterozygous diploid where only one of the *VMA1* alleles carries PI-Sce1, the endonuclease cleaves the *VMA1* sequence lacking the endonuclease-coding portion to initiate homing, which consists of a recombination reaction associated or not with a CO (Figure S7). Using *de novo* genome assembly, we defined which version of *VMA1* was carried by each parental strain as well as in the other 26 *L. kluyveri* natural genomes already sequenced [23]. In total, nine strains harbor a *VMA1* gene with PI-Sce1, including our parental strain NRBC10955, and nineteen isolates harbor an empty *VMA1* gene, including the other parental strain 67-588. Therefore, meiosis specific homing of PI-Sce1 explains the high recombinogenic activity at the *VMA1* locus observed in our dataset that includes five COs, five double COs, nine short 4:0 NCOs and ten LOH events (Figure S6). Note that in a single meiosis, PI-Sce1 can cut one or two chromatids generating 3:1 and 4:0 NCOs as well as single and double COs, respectively. Therefore, in this specific case, only borders of long tracts of LOH can be unambiguously attributed to RTG events.

### *L. kluyveri* has a lower recombination rate than *S. cerevisiae*

To estimate the number of recombination events per meiosis in *L. kluyveri,* we considered independently double COs, 4:0 NCOs and large LOHs that are considered to result from RTG events. This results in 334 COs without GC, 541 single COs with 3:1 GC tracts (Figure 1 and S5) and 274 NCOs in 49 meioses. We also observed that at least 100 single COs occurred within LOH regions (see below). This corresponds to 19.9 COs and 5.6 NCOs per meiosis on average. RTG events comprise 138 double COs, 178 large LOH tracts within chromosomes, 114 large LOH tracts encompassing one telomere and 132 4:0 NCOs. Each double CO corresponds to one CO per RTG. Large internal LOH tracts account for two COs per RTG. Telomere proximal LOH tracts account for one CO per RTG. Altogether, we observed 608 COs and 132 NCOs within 38 meioses, corresponding to 16 COs and 3.5 NCOs per RTG on average considering that only one RTG occurred.

The detected recombination events per RTG correspond to an underestimation of the actual number of events since we analyzed only one of the two daughter cells after RTG (Figure S5). Considering that not all of the recombination events occurring during RTG are detected (as describe in [31]), we estimate that the average number of COs per RTG is similar to those per meiosis (Figure S8).

The presence of LOH regions also leads to an underestimation of meiotic recombination events that occur in such regions. Even numbers of COs within a LOH region involving only two chromosomes are missed as well as all NCOs. Considering the global recombination rate, we estimate that about 190 COs are expected within LOH regions, 90 of which being not undetectable (see Methods).

Both COs and NCOs are significantly less abundant than in *S. cerevisiae* that has a similar genome size, and where 73 to 90 COs and 27 to 46 NCOs have been detected per meiosis, depending on the studied hybrid [6,24]. We used a stringent threshold for the detection of conversions (see Methods). This may have led us to miss some short conversions. Indeed, the median size of *L. kluyveri* conversion tracts is 2.9 kb for those associated with COs, and 2.2 kb for NCO conversion tracts (Figure 1B) compared to 2 kb and 1.8 kb in *S. cerevisiae,* respectively [6]. However, our strategy should reveal virtually all COs. Therefore, we can conclude that the genetic map of the *L. kluyveri* hybrid used is at least four times smaller than that of *S. cerevisiae.*

As previously observed in *S. cerevisiae* and in other species, the average number of COs per chromosome is correlated to chromosome length in *L. kluyveri*. Remarkably, the linear relation between chromosome size and number of COs displays a positive intercept of 1.0 CO for a virtual chromosome size of 0 bp both in *S. cerevisiae* and *L. kluyveri* (Figure 3A). This suggests that in these species the meiotic program is set up in such a way that at least one CO will form per chromosome independently of its length.

**Figure 3.**
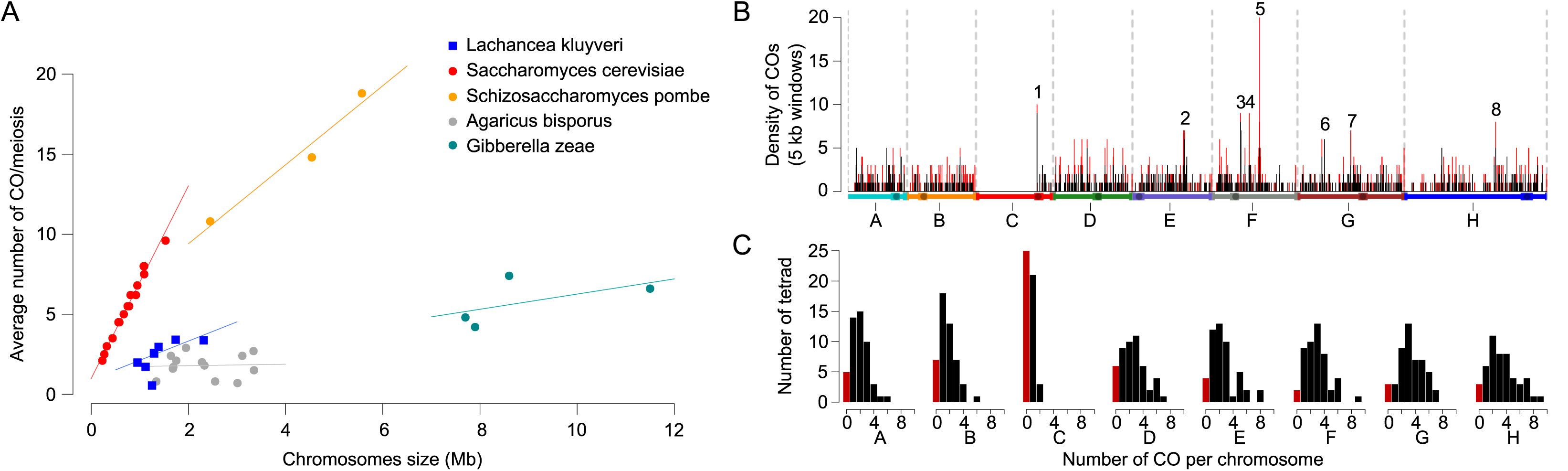
Recombination landscape in *L. kluyveri.* (A) Average number of COs per meiosis per chromosome for five species: Saccharomycontina (*Lk:* intercept = 0.92, slope = 1.2, *Sc:* intercept = 0.99, slope = 6.0), Taphrinomicotina (*Sp:* intercept = 4.5, slope = 2.5), Agaricomycotina (*Ab:* intercept = 1.7, slope = 0.05), Pezizomycotina (*Gz*: intercept = 1.5, slope = 0.47). Data obtained from [34–38]. (B) Density of COs along the genome using a 5 kb window. Regions 1 to 8 correspond to hotspots of COs. Black peaks correspond only to single COs. Red peaks included pre-RTG events (double COs and LOHips). (C) Histogram of the number of COs per meiosis for each *L. kluyveri* chromosome. The red bars correspond to the cases where no CO was detected on the chromosome.

*S. cerevisiae* and *L. kluyveri* have a similar genome size of approximately 12 Mb and therefore the recombination rate is estimated to ~1.6 COs/Mb and ~6 COs/Mb for *L. kluyveri* and *S. cerevisiae,* respectively. This difference underlines the particularly high recombination rate of *S. cerevisiae* that is rather uncommon in other eukaryotes (Figure 3A). Indeed, the fission yeast *Schizosaccharomyces pombe,* which also harbors a 12.6 Mb genome size, displays a number of COs ranging from 34 to 44 per meiosis, depending on the hybrid studied, leading to approximately 3 COs/Mb per meiosis [34,35]. Recombination analyses were also performed in other more distant fungi species such as *Agaricus bisporus* and *Gibberella zeae,* where the recombination was estimated to be ~0.78 COs/Mb and ~0.65 COs/Mb per meiosis, respectively [36–38].

The size of the smallest chromosome might explain part of the difference in recombination rates [39]. Indeed, species carrying small chromosomes tend to display high recombination rates to stochastically ensure at least one CO on the smallest chromosome. This is illustrated by a negative correlation between the size of the smallest chromosome and the overall recombination rate across species [39]. We obtained meiotic data from 30 species to complete the one obtained in *L. kluyveri* and confronted their recombination rates to the size of their smallest chromosome. As expected, we observed a negative correlation between these two parameters (R^2^ = 0.79, Figure S9) and *L. kluyveri* recombination data are consistent with this correlation. Whether the chromosome number, 8 *vs.* 16 in *L. kluyveri* and *S. cerevisiae,* respectively, impacts the global meiotic recombination rate is unknown.

Mechanistically, several hypotheses could explain the reduced rate of COs in *L. kluyveri,* such as less initiating meiotic DSBs than in *S. cerevisiae,* or a more frequent use of the sister chromatid as a template for DSB repair. We therefore compared the DSB level between *S. cerevisiae* and *L. kluyveri* on chromosomes of similar length by pulse-field gel electrophoresis (PFGE). We performed this experiment in the homozygous diploid CBS10367 isolate that has been independently chosen for its fast and efficient sporulation properties, although it shows a two-hour delay in meiotic entry compared to the *S. cerevisiae* SK1 isolate (Figure S10A). At comparable time points after meiotic induction, *i.e.* four and six hours, respectively, we found similar DSB levels, suggesting that the four-fold lower level of recombination in *L. kluyveri* does not come from a corresponding decrease in initiating DSBs (Figure S10B).

### The 1-Mb Sakl0C-left region is associated with a low recombination activity

Chromosome C stands as an outlier with significantly less COs per meiosis compared to the other chromosomes (Figure 3A). This specifically comes from the low recombination activity of the 1-Mb left arm of the chromosome C (Sakl0C-left) where no CO was observed and only one NCO in the *PAC10* gene was detected (Figure S6). *De novo* genome assemblies of the two parental genomes using mate pair sequencing data (see Methods) show that the two chromosomes C are collinear to the chromosome C of the CBS3082 reference strain (Figure S11). Consequently, the absence of CO on Sakl0C-left is not due to the presence of chromosomal rearrangements such as inversions that could prevent recombination. This low recombination activity of the Sakl0C-left region is unexpected and adds up to the other peculiarities of this genomic region. We recently performed a *L. kluyveri* population genomic survey and showed that this specific region is a relic of an introgression event, which occurred in the last common ancestor [23]. Our study also revealed that the Sakl0C-left region underwent a molecular evolution pattern different from the rest of the genome. It is characterized by a higher GC content, a higher sequence diversity within isolates (π = 0.019 *vs.* 0.017) as well as a dramatically elevated frequency of G:C → A:T substitutions. Therefore, the higher density of mutations on Sakl0C-left between our parental strains compared to the rest of the genome (1.13% *vs.* 0.69%) could prevent the use of the homologous chromatid in DSB repair because of the antirecombination activity of the mismatch repair machinery [24,40]. Another possibility is that fewer DSBs are induced in this region, this being potentially related to the high GC content. To test this latter possibility, we monitored the DSB level on several chromosomes including chromosome C by Southern blot after PFGE of samples from meiotic time courses using the CBS10367 isolate (Figure 4). We observed DSB hotspot signals all along both chromosomes D and E. Only one centromere proximal hotspot was detected on Sakl0C-left. Finer mapping of this DSB hotspot showed it is located in the promoter region of *GPI18,* matching the CO hotspot number 1 (Figure S12 and see below). This result is remarkable since it reveals a 1Mb region deprived of detectable DSBs. Interestingly, even if the *GPI18* hotspot is located in the Sakl0C-left, it lies close to the centromere outside but flanking the GC rich region. The next unanswered challenge is to determine what makes such a large region refractory to Spo11 cleavage.

**Figure 4.**
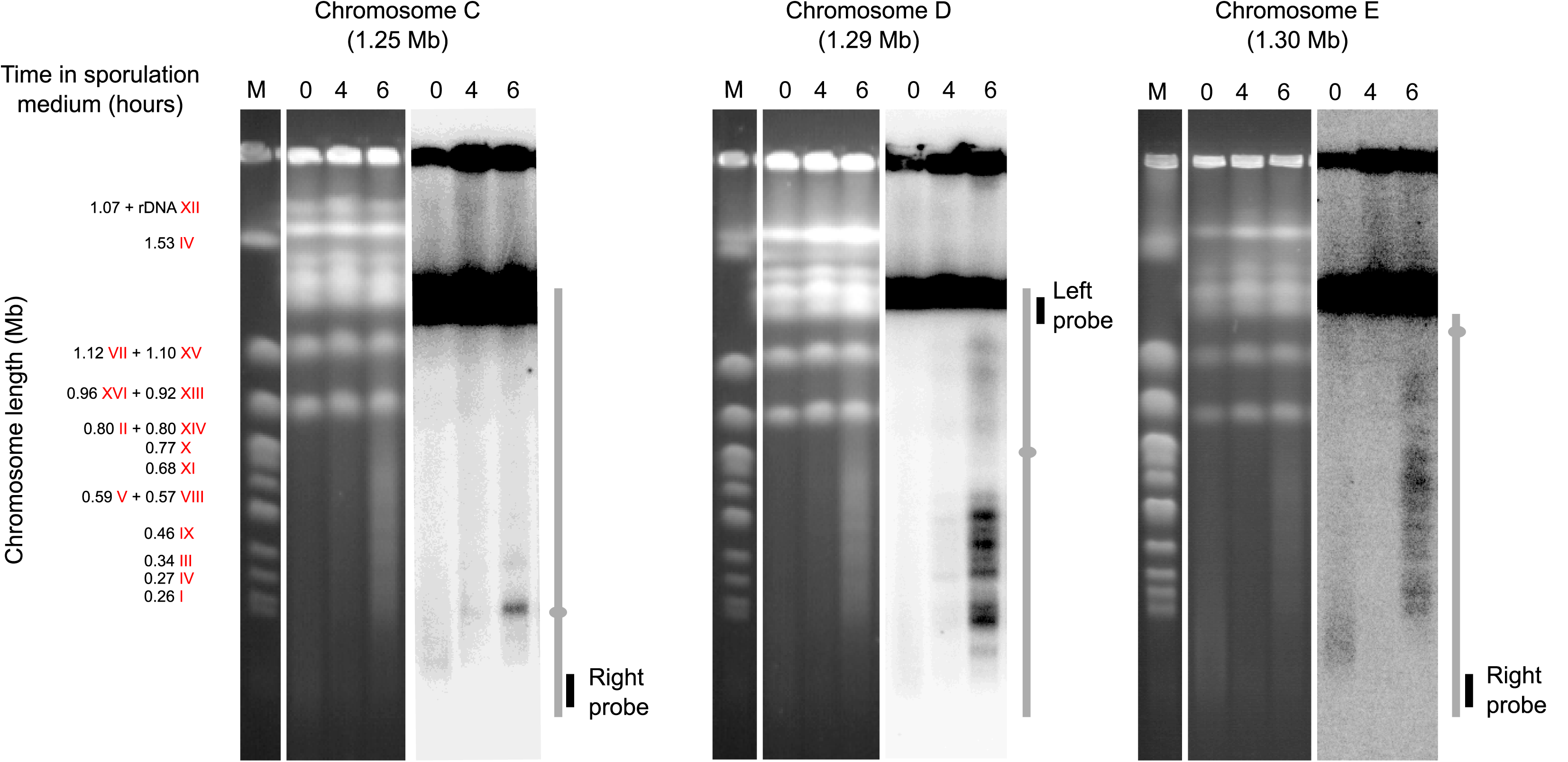
Visualization of meiotic DSB hotspots on *L. kluyveri* chromosomes C, D and E. *L. kluyveri* meiotic chromosomes were separated by PFGE and revealed with telomere proximal probes after Southern blot. The strain CBS10367 *Δsae2/Δsae2* was used for a better visualization of the broken chromosomes. Left parts, ethidium bromide stained gels. Right parts, radioactive signals of the corresponding membranes after Southern blot. M, molecular size marker corresponding to mitotic *S. cerevisiae* chromosomes. Chromosomes are represented vertically, with the location of the probe used. Grey dots indicate centromeres location. Note that the exposure of chromosome C was increased compared to chromosome D and E to better visualize the centromeric proximal DSB hotspot. This underlines the weaker DSB level on chromosome C.

Intriguingly, we previously observed that the Sakl0C-left region has an increased ancestral recombination rate estimate ρ (based on linkage disequilibrium decay) compared to the rest of the genome (ρ = 8.5 *vs.* 3.2 Morgans/kb, respectively) [23]. This observation stands in contrast to the actual recombination rate we determined here. Several reasons could explain this discrepancy among which an evolution of the rate over time [23,41]. More generally, the Sakl0C-left region reveals an unusual genetic behavior at the scale of a chromosome. Interestingly, the *MAT* locus is located in this region and the absence of recombination reminds of its suppression associated to the *MAT* locus in fungi species such as *Neurospora tetrasperma* or *Cryptococcus* spp. [42,43], or more generally to sex chromosomes such as the chromosome Y in mammals. The reason of this suppression is unclear but the lack of recombination might result in a long-term degeneration of genes [44].

### High incidence of non-recombining chromosomes in *L. kluyveri* tetrads

A consequence of the absence of recombination in the Sakl0C-left is a global deficit of COs on chromosome C (Figure 3A). Strikingly, 25 out of 49 tetrads show no CO on chromosome C but correct chromosome segregation. In addition, more than half of the *L. kluyveri* tetrads on average contained another chromosome without any CO (E0, for 0 exchange) while it happened in only one tetrad out of 46 in *S. cerevisiae* [6] (Figure 3C). Since the presence of E0 chromosomes threatens accurate chromosome segregation at meiosis I, this result suggests either inaccurate chromosome segregation in *L. kluyveri,* notably for chromosome C, and therefore elevated spore death, or the existence of an efficient compensatory mechanism such as distributive segregation [45]. We could not determine the spore viability of the parental strains of our hybrid because no corresponding homozygous diploid was available. However, we determined that the spore viability of the *L. kluyveri* reference strain CBS3082 is close to 100% showing that at least this isolate undergoes proper homolog segregation at meiosis I. Assuming the high frequency of E0 is a general property of this species, this tends to support the existence of an efficient compensatory mechanism for meiotic chromosome segregation in *L. kluyveri*.

### CO interference in *L. kluyveri*

By analyzing inter CO distances, we could determine if *L. kluyveri* CO interfere. CO interference is a phenomenon that tends to ensure an even distribution of COs along chromosomes and that may promote efficient chromosome segregation at meiosis I. While the mechanism of CO interference is still unclear, the so-called ZMM group of proteins Zip1, 2, 3, 4, Spo16, Msh4, 5 and Mer3 promote CO interference in *S. cerevisiae* [46]. By contrast to the other *Lachancea* species, *L. kluyveri* conserved all these proteins [47], which led us anticipate the presence of CO interference. We looked for CO interference by measuring inter-CO distances and fitting them to a gamma distribution function characterized by a shape (*k*, estimating interference) and scale (*θ*, estimating the inverse of CO rate) parameter [25,48,49]. We first confirmed that *L. kluyveri* displayed CO interference by comparing inter-CO distances distribution to a gamma distribution with no interference (shape parameter (*k*) = 1, Kolmogorov-Smirnov test *P* = 0.0072). However, the optimal estimation of the strength of interference in *L. kluyveri* is significantly lower than that of *S. cerevisiae* (*k* = 1.47 *vs.* 1.96, Kolmogorov-Smirnov test *P* = 0.015). Additionally, because the COs frequency is low, the scale parameter in *L. kluyveri* is higher than in *S. cerevisiae* (*θ* = 187 *vs.* 61.7) (Figure S13A). Remarkably, the distribution of size of LOH region follows approximately the same gamma law (*k* = 1.45 and *θ* = 213, Figure S13B), which is in agreement with their meiotic origin.

### Recombination hotspots are not conserved between *L. kluyveri* and *S. cerevisiae*

To identify CO hotspots (Figure 3B and Figure S6), we determined the density of COs along the genome using a 5 kb window, and defined as hotspots the regions displaying seven or more COs per 5 kb (*P* < 0.02 based on 10^5^ randomly generated COs positions). In addition to the *VMA1* hotspot discussed previously, seven other CO hotspots were identified across the 49 meioses and 38 RTGs. They are located in the *GPI18, RAS1, PIS1, PRM2, OST6, VMA10,* and *ALT2* orthologs (hotspot 1, 2, 3, 4, 6, 7, and 8, respectively) (Figure 3B).

As mentioned previously, the *GPI18* CO hotspot is on the left arm of chromosome C at approximately 9 kb from the centromere (hotspot 1, Figure 3B). This corresponds exactly to the end of the 1-Mb introgressed region (Sakl0C-left) and carries relics of transposable elements in both parental strains.

Whether these transposable elements influence DSB formation or not remains to be established, since such the presence of such elements is not predictive of the local recombination activity [50]. Another hypothesis is that we enriched the instance of COs happening on the chromosome C by selecting only full viable tetrads, as the absence of recombination on the Sakl0C-left region might lead to a reduction of the spore viability. In any case, this CO hotspot correlates with the DSB hotspot we observed in the promoter region of *GPI18* from the CBS10367 isolate that also has relics of transposable elements (Figure S12).

The six remaining CO hotspots are associated with genes that mostly display a high expression level (above the median) during growth in rich medium (Table S1) [41]. As for *GPI18,* we observed corresponding DSB hotspots in the promoters of *RAS1* (chromosome E) and *PIS1* (chromosome F) (Figure S12). Remarkably, none of the well-known recombination hotspots in *S. cerevisiae* were significantly detected in *L. kluyveri (e.g. IMG1, HIS4, CYS3)* [6,8]. In addition, among the eight hotspots detected in *L. kluyveri,* only one is significantly enriched for COs in *S. cerevisiae* considering the meiotic recombination dataset from Mancera *et al.* [6]. We specifically tested the significance of the reduction of the number of proximal COs in *S. cerevisiae* for each couple of orthologs involved in *L. kluyveri* [6]. We confirmed the difference for all the recombination hotspots with the exception of the one located in the syntenic block *MSC7-VMA10-BCD1* (using a 10 kb window, Fisher exact test: P < 0.01). At a larger scale, we compared between the two species the number of proximal COs for all couples of orthologs and observed a poor correlation (R^2^ < 0.01, Figure 5). The *VMA10* hotspot appears to be the only one conserved between the two species. Interestingly, the *Saccharomyces* species (e.g. *S. cerevisiae, Saccharomyces paradoxus*), which are closely related, display similar or conserved recombination patterns, as shown by using population genomics data and comparing genome-wide recombination initiation maps [15,17]. Here, our results demonstrate that the distribution of meiotic recombination is not persistent across distantly related yeast species covering a large evolutionary scale, exceeding the one of the *Saccharomyces* genus. The most probable factor explaining the conservation of repartition of recombination across this genus is likely the shared synteny and chromosomal organization.

**Figure 5.**
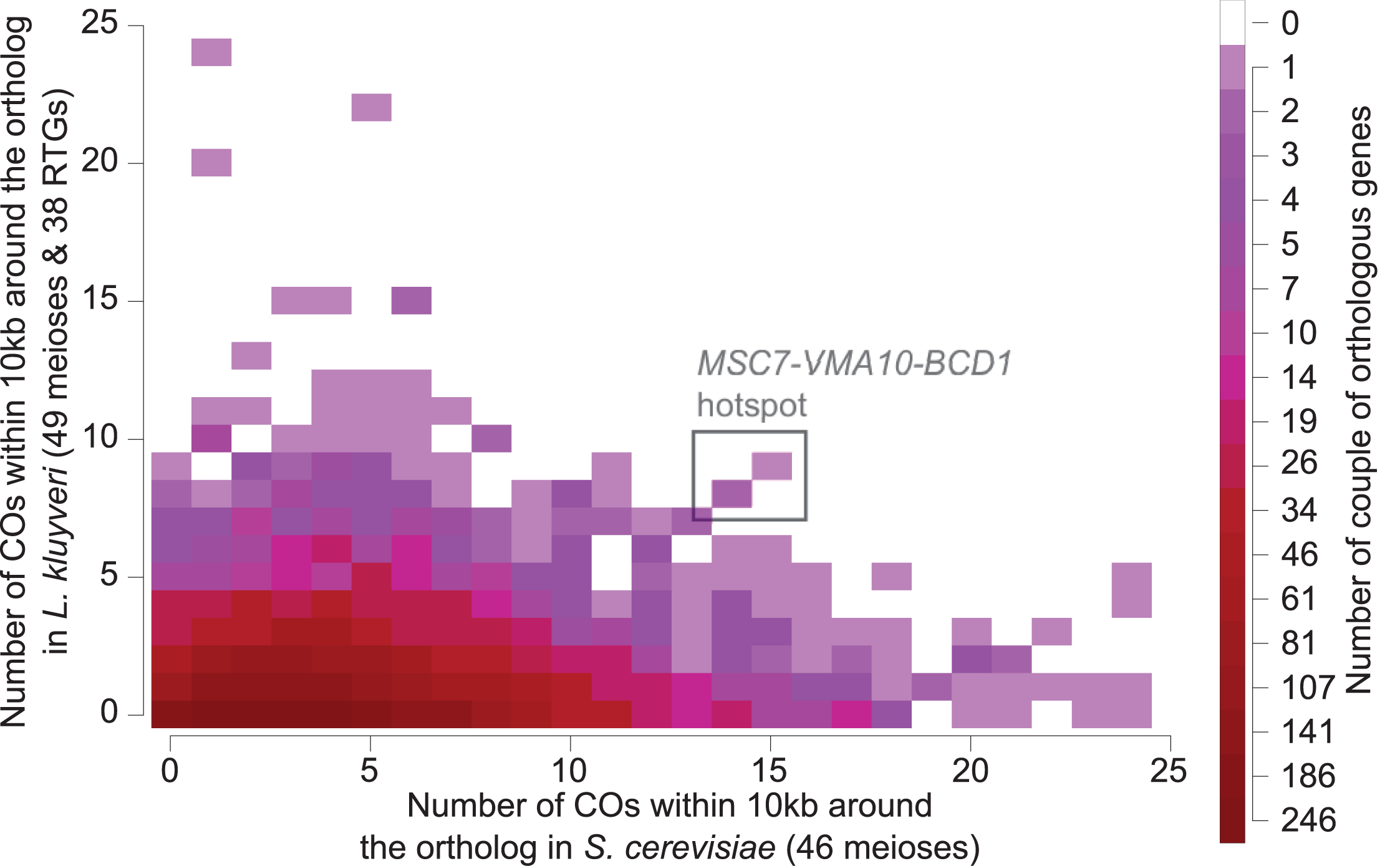
Comparison of the number of proximal COs between *S. cerevisiae* and *L. kluyveri* orthologs. For each ortholog pair, we determined the number of COs in a 10kb window around each gene. The plot displays the density of ortholog pairs according to the associated number of proximal COs. The conservation of COs position between species would generate a linear correlation, with several instances located in the upper-right part of the plot. Here, the R^2^ of the linear regression is 1.16× 10^-3^ (P = 8.22× 10^-3^). The unique common hotspot between the two species is *MSC7-VMA10-BCD1.*

## Conclusion

Genome evolution is strongly tied to the meiotic behavior and recombination landscape. With the exploration of the meiotic genetic exchanges of a protoploid yeast species and its comparison to *S. cerevisiae* as well as other fungi (Figure 3A), we unveiled characteristics that bring a new light on the variation of the recombination landscape and its potential impact on evolution. The lack of recombination hotspot conservation between *L. kluyveri* and *S. cerevisiae* provides new insight into the understanding of the biological mechanism controlling these hotspots as it could be associated to the differences in synteny.

A striking feature of *L. kluyveri* is its low recombination rate in a 1-Mb region of the chromosome C, showing an absence of CO for about half of the surveyed tetrads. We showed that this results from a lack of Spo11-induced DSB in the corresponding region. What prevents Spo11 from making DSBs over such a long region and the consequences of this meiotic defect remain to be determined. Interestingly, the absence of recombination in this introgressed region that determines mating-types has the consequence to genetically link all the genes and therefore all phenotypic characteristics their alleles might induce, which potentially hindered species diversification and adaptation.

The capacity for *L. kluyveri* hybrids to undergo RTG during stress, resulting in a high number of LOH regions, is an unexpected source of genetic and therefore phenotypic diversity. Meiotic reversions are generally not taken into account in the exploration of species evolution. Here, our data show that the control of commitment into meiosis might be weaker in some yeast species, increasing allelic shuffling during stress. The significance of this evolutive advantage needs to be further investigated as it has been underestimated so far.

Our unexpected observations obtained by the deep exploration of recombination landscape of a non-conventional yeast species also demonstrate that yet many mechanisms and variations affecting evolution still remain to be described and explored.

## Materials and Methods

### Strains construction, growth conditions and tetrad dissection

Both parental strains NBRC10955a (*MATa*) and 67-588 (*MATα*) had their *CHS3* gene encoding for chitin synthase inactivated to facilitate tetrad dissection [23,51]. Cells were electroporated with a DNA fragment containing the drug resistance marker *KanMX,* produced by fusion PCR [29]. Gene replacement was validated by PCR. All primers are listed in table S2.

Growth and cross were performed on YPD (yeast extract 1%, peptone 2%, glucose 2%) at 30°C. Sporulation was induced by plating cells on sporulation medium (agar 2%, potassium acetate 1%) and incubating 2 to 7 days at 30°C. The asci of the tetrads were digested using zymolyase (0.5 mg/mL MP Biomedicals MT ImmunO 20T) and the spores were isolated using the MSM 400 dissection microscope (Singer instrument). The whole genome of each of the 196 spores from the 49 full viable tetrads selected was sequenced.

### Genomic DNA extraction, sequencing, SNP calling and data analysis

Genomic DNA was extracted using the MasterPure Yeast DNA Purification Kit (tebu-bio) and sequenced using Illumina HiSeq 2000 technology. We used paired-end libraries and 100 bp per read. The average reads coverage is 204x and 100x for NBRC10955a and 67-588, respectively, and around 70x for each of the 196 spores. The reads obtained were cleaned, trimmed and aligned to the reference genome using BWA (-n 8-o 2 option). Raw data are available on the European Nucleotide Archive (http://www.ebi.ac.uk/ena) under accession number PRJEB13706.

To select SNPs to be used as genotyping markers, the reads of the parental strains NBRC10955a and 67-588 were mapped against the reference genome, in which repetitive and transposable elements were masked with RepeatMasker. SAMtools was used to call variants, and a set of high quality differential SNPs between parental strains was generated. To be integrated in this set, differential positions must be, for both parents, (I) covered between 50x and 300x, (ii) have a mapping quality score higher or equal to 100, (iii) have an associated allele frequency (AF tag in VCF file) of 0 or 1, and (iv) should not be located in close proximity (10 base pairs) to an indel (insertion/deletion). The final set consists of 58,256 markers used to genotype the progeny.

The reads associated with each spore were mapped and variants called with the same strategy as for the parental strains. At each position of the set, a parental origin was assigned if the residue (i) corresponds to one parental version, (ii) shows a coverage higher than 25% of the average coverage of the spore (20x in most cases), (iii) has a mapping quality higher than 100, and (iv) displays an allele frequency of 0 or 1. If not, a NA flag was assigned to the position.

From this matrix (58,256 × 196 genotypes), we discarded markers flagged NA across 60 spores (30%) or more, markers displaying unbalanced genotype origin in the population (fraction of parental origins above 70% / under 30%) as well as subtelomeric regions carrying translocated and duplicated regions.

Finally 56,612 markers corresponding to high quality differential SNPs between parents and for which most of the spores displayed a reliable allelic origin were used.

**Identification of recombination events**

To identify the localization and size of recombination events, including GCs with or without COs and LOH, we adapted the CrossOver python scripts from the Recombine pipeline (http://sourceforge.net/projects/recombine/) [25] to fit *L. kluyveri* genomic characteristics. For each tetrad, we disregarded markers for which at least one spore displayed a NA flag before running CrossOver. In order to conserve only reliable information, a GC event was considered only when at least three markers were involved, which corresponds to 63% of the detected GCs. We defined a LOH as a 4:0 GC event larger than 10 kb (Figure 1). LOH regions detected were validated in one tetrad by looking at the sequence data directly. To do so, pooled reads of all four spores from the selected tetrad were aligned to the reference genome (BWA), and the polymorphic positions were identified (SAMtools). Regions without LOH would present both parental allelic versions and thus appear to be heterozygous, whereas LOH regions would be homozygous. The detected homozygous regions coincide with LOH regions identified using CrossOver (Figure S4). All detected meiotic recombination events are provided in table S3.

The number of COs occuring within LOH regions was estimated using the recombination rate. A total of 875 COs has been detected on the genomes of 49 meioses with the exclusion of the Sakl0C-left region and LOH regions (416.4 Mb). The cumulative size of all the LOH regions was 90.4 Mb. With the hypothesis that the recombination rate is homogeneous (with the exclusion of Sakl0C-left region), we estimated that 190 COs occurred within LOH regions of our meiotic sample.

### Quantification of meiotic DSBs

*S. cerevisiae* SK1 [52] and *L. kluyveri* CBS10367 [23] diploid strains homozygous for the *sae2* null allele were used to detect the level of cut chromosomes during meiotic time courses. Procedures relative to *S. cerevisae,* Southern blot analysis and chromosome preparation in agarose plugs are as described in [52]. Oligonucleotide sequences for probe synthesis are in table S2. *L. kluyveri* cells were inoculated in pre-sporulation medium (0.5% yeast extract, 1% peptone, 0.17% yeast nitrogen base, 1% potassium acetate, 1% ammonium sulfate, 0.5% potassium phthalate, pH 5.5) over night at 30°C to reach an OD_600_ of 1, washed once with water and resuspended in 1% potassium acetate at 30°C. Cells were harvested at different time points, washed with water and frozen. Pulse field gel electrophoreses were performed using a Bio Rad CHEF DRIII apparatus at 14°C, initial switch time 60 seconds, final switch time 120 seconds, 6 V/cm, angle 120° for 20h.

### *de novo* assembly of the parental genomes

SOAPdenovo (v2.04, [53]) was used to construct *de novo* assemblies for both parental strains using a combination of paired-end and mate-pair Illumina reads (European Nucleotide Archive: PRJEB13706). We tested a range of k-mer size (from 45 to 83) and compared the obtained assemblies at the contiguity level. We found k=63 and k=71 to be optimal for the 67-588 and NBRC10955 parental strains, respectively. The selected assemblies were compared to the reference genome with nucmer [54] and results were plotted with mummerplot (Figure S11).

## Acknowledgments and funding information

We are grateful to Gilles Fischer for fruitful discussions and his invaluable advice, and to Marie-Pauline Beugin and Kenny Dubois for *L. kluyveri* strain constructions and meiotic protocols set up. This work was supported by grants from the Agence Nationale de la Recherche (ANR): grant 2013-13-BSV6-0012-01 to BL, and grant ANR-16-CE12-0019 to JS. We also thank the University of Strasbourg Institute for Advanced Study (USIAS) for their financial support. JS is a member of the Institut Universitaire de France.

